# The gliotransmitter S100β regulates synaptic plasticity in the visual cortex

**DOI:** 10.1101/2025.07.29.667525

**Authors:** Yanis Inglebert, Rafael Sanz-Gálvez, Arlette Kolta

**Affiliations:** Department of Neurosciences, Université de Montréal, Montréal, Canada; Centre Interdisciplinaire de recherche sur le cerveau et l’apprentissage (CIRCA), Montréal, Québec, Canada; Faculté de Médecine Dentaire, Université de Montréal, Montréal, Québec, Canada

**Keywords:** Astrocytes, Synaptic Plasticity, STDP, S100β, Calcium

## Abstract

Synaptic plasticity is a fundamental mechanism of memory storage in the brain. Among the various rules governing changes in synaptic strength, Spike Timing-Dependent Plasticity (STDP) stands out for its strong physiological relevance *in vivo*. Ubiquitous across brain regions and neuronal types, STDP is a complex and multifactorial process influenced by factors such as neuromodulation, extracellular calcium levels, and activity patterns. However, one relatively understudied factor is the role of astrocytes, despite their well-established involvement in regulating synaptic transmission and neuronal excitability through gliotransmitter release. While some factors have garnered significant attention, others, like S100β, have remained relatively underexplored despite their potential importance in regulating synaptic plasticity. S100β is a calcium-binding protein, allowing it to influence extracellular Ca²⁺ concentration and potentially all Ca^2+^-dependent plasticity processes. Building on our previous research in the visual cortex, where we examined the regulation of neuronal excitability by S100β, we chose to further investigate the role of astrocytes and S100β in synaptic plasticity at layer 2/3-layer 5 synapses in the visual cortex. We demonstrated that S100β is an important gliotransmitter to consider, capable of regulating long-term potentiation.

## INTRODUCTION

Spike Timing-Dependent Plasticity (STDP) is currently the leading theoretical framework for understanding synaptic plasticity and how the brain processes and stores information (Debanne and Inglebert, 2023). Observed both *in vivo* and *in vitro* across various species, including humans, STDP relies on the precise timing of pre- and post-synaptic activity to encode information (Feldman, 2012). Traditionally, long-term potentiation (t-LTP) occurs when presynaptic activity, in the form of excitatory postsynaptic potentials (EPSPs), precedes postsynaptic action potentials. Conversely, long-term depression (t-LTD) arises when presynaptic activity follows postsynaptic activity (Debanne et al., 1996; Bi and Poo, 1998). Although STDP is a universal plasticity rule, it is a complex and multifactorial process which expression varies across synapses, depends on distinct mechanisms, and is influenced by different regulatory factors such as activity patterns (Pike et al., 1999), neuromodulation (Brzosko et al., 2019), and extracellular calcium concentrations ([Ca^2+^]_e_) (Inglebert et al., 2020; Inglebert and Debanne, 2021). In some cases, it is gated by specific neuromodulators (Zhang et al., 2009; Huang et al., 2013), such as monoamines, while in others, particular activity patterns, such as specific frequencies or bursts, are necessary for its induction (Pike et al., 1999; Inglebert et al., 2020). This diversity makes STDP a highly heterogeneous mechanism, requiring careful examination across different synapses to build a comprehensive framework of the rules governing synaptic modifications (McFarlan et al., 2023). Among factors susceptible to influence STDP, astrocytes have largely been ignored despite being known for regulating synaptic transmission and neuronal excitability (Araque et al., 1998, 2014; Henneberger et al., 2010; Tan et al., 2017). To date only a few studies have investigated the role of astrocytes in STDP (Min and Nevian, 2012; Pérez-Rodríguez et al., 2019; Falcón-Moya et al., 2020; Martínez-Gallego et al., 2022; Sanz-Gálvez et al., 2024), and those examining their effects on synaptic plasticity focused primarily on gliotransmitters such as glutamate, adenosine, or D-serine, which have been described to gate t-LTP or t-LTD (Min and Nevian, 2012; Pérez-Rodríguez et al., 2019; Falcón-Moya et al., 2020). However, little attention has been given to S100β, a calcium-binding protein released from astrocytes that belongs to the S100 family (Donato et al., 2013). Our previous work in the brainstem and visual cortex has shown that S100β secretion lowers [Ca^2+^]_e_, leading to changes in intrinsic excitability and neuronal firing patterns (Morquette et al., 2015; Ryczko et al., 2021; Gaudel et al., 2025). STDP is highly sensitive to [Ca²⁺]ₑ, and studies in both the hippocampus and cortex have shown that investigating STDP under reduced [Ca²⁺]ₑ can produce results opposite to those observed under higher calcium conditions (Inglebert et al., 2020; Inglebert and Debanne, 2021; Chindemi et al., 2022). This implies that S100β may regulate or modulate synaptic plasticity levels or thresholds by altering the local Ca^2+^ environment. To explore its potential regulatory role, we investigated t-LTP at synapses from layer 2/3 to layer 5 (L2/3-L5) in the visual cortex.

## RESULTS

### Extracellular calcium is a strong regulator of t-LTP

First, to assess whether [Ca^2+^]_e_ affects t-LTP at L2/3-L5 synapses of the visual cortex as in the hippocampus and in predictions from computational models, we induced t-LTP under physiological [Ca^2+^]_e_ (1.2 mM) and under a [Ca^2+^]_e_ commonly used in *in vitro* studies (2 mM), in acute slices obtained from P14-P21 wild-type (WT) mice. The t-LTP induction protocol involved a single presynaptic stimulation in the L2/3 followed 10ms later by a single postsynaptic action potential elicited by current pulse injection through the recording electrode in an L5 pyramidal neuron (L5PN), repeated 100 times at varying frequencies (1, 10, 20 and 50 Hz) to capture the full spectrum of t-LTP (**Fig 1A**). At 2 mM, a robust t-LTP is induced at 10 Hz (133.1 % ± 19.2, n = 9, **Fig 1C**), 20 Hz (137 % ± 13.3, n = 3, **Fig 1D**) and 50 Hz (132 % ± 18.2, n = 7, **Fig 1E**) while, in average, no change is observed at 1 Hz (101.4 % ± 14, n = 8, **Fig 1B**). Lowering the [Ca^2+^]_e_ to 1.2 mM led to a similar t-LTP at 50 Hz (128.2 % ± 15.3, n = 7, **Fig 1D**, but to drastically different outcomes, shifting t-LTP to t-LTD at 10 Hz (76.8 % ± 10.5, n = 6, **Fig 1C**) and 20 Hz (83.5 % ± 6.2 n = 6, **Fig 1D**). In addition, at 1 Hz while no plasticity was observed at 2 mM, the [Ca^2+^]_e_ reduction led to a robust t-LTD (75.5 % ± 11.7, n = 10, **Fig 1B**). Together, these findings suggest that [Ca2+]_e_ is a key regulator of synaptic plasticity at L2/3-L5 synapses in the visual cortex and support the hypothesis that astrocytes may influence synaptic plasticity through secretion of S100β. In the following experiments, we focused on the 10 Hz frequency, as it was sufficient to elicit robust t-LTP and t-LTD depending on the [Ca2+]_e_. The hypothesis being that greater release of S100β should lead to a decrease in [Ca2+]_e_ as previously shown, a shift of t-LTP to t-LTD in 2 mM [Ca2+]_e_, while blockade of S100β should shift t-LTD to t-LTP in 1.2 mM [Ca2+]_e_.

**Figure 1.**
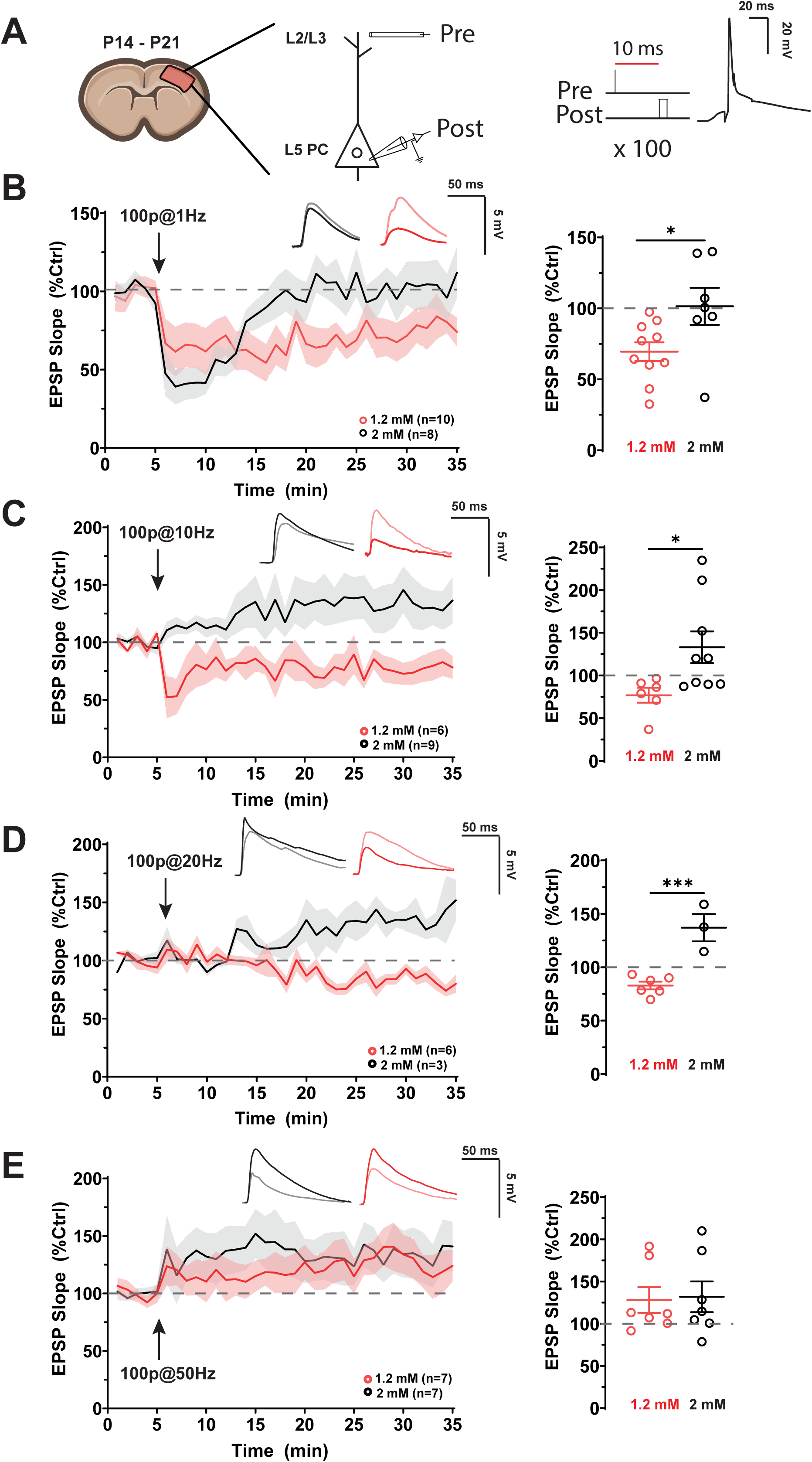
Extracellular calcium regulates t-LTP at L2/L3-L5 synapses in the visual cortex. **A.** Experimental paradigm. Whole-cell recordings were obtained from layer 5 pyramidal neurons in acute coronal slices, while a stimulating pipette was placed in layer 2/3 to evoke presynaptic stimulation. The STDP protocol consisted of 100 pre-post pairings repeated at different frequencies. **B,C,D and E.** *Left:* Time course of normalized EPSP slope before (light traces) and after (dark traces) pre-post stimulation at 1, 10, 20, or 50 Hz in two different extracellular calcium concentrations: 2 mM (black) and 1.2 mM (red). *Right:* Quantification of EPSP slope changes in the last 10 minutes of recording, with individual values for each cell represented as red open circles (1.2 mM) and black open circles (2 mM).

### Astrocytes activity can influence t-LTP at L2/3-L5 synapses

First, to determine whether astrocytic activity modulate the plastic changes observed at each [Ca2+]_e_, we inactivated the peri-neuronal astrocytic network using dual-cell recordings of a L5PN and a neighboring astrocyte located near its soma (47.4 µm ± 11.3, **Fig 2B Right;** 45 µm ± 10.4. **Fig 2C, Right**). The astrocyte was filled with BAPTA, a calcium chelator commonly used to suppress astrocyte activity (**Fig 2A**). Astrocytes were visualized using SR-101 after pre-incubating the slices for 5 minutes at room temperature, as described in Watanabe et al., 2023. Interestingly, all plastic changes were abolished in this condition (101.2 % ± 4.8, n = 7, in 2mM Ca^2+,^ **Fig 2B**, and 99,9 ± 17, n=6, in 1.2mM Ca2+, **Fig 2C**). Notably, since astrocytes were also filled with Alexa 488, we could visualize the extent of the astrocytic network that was inactivated by the end of the recordings, as indicated by the white arrows in **Figure 2A**. These findings suggest that peri-neuronal astrocyte activity is essential for the normal expression of t-LTP or t-LTD at L2/3-L5 synapses in the visual cortex. Our data indicate that peri-neuronal astrocyte activity can modulate t-LTP. Altogether, our results highlight astrocytes as key partners in t-LTP at L2/3-L5 synapses.

**Figure 2.**
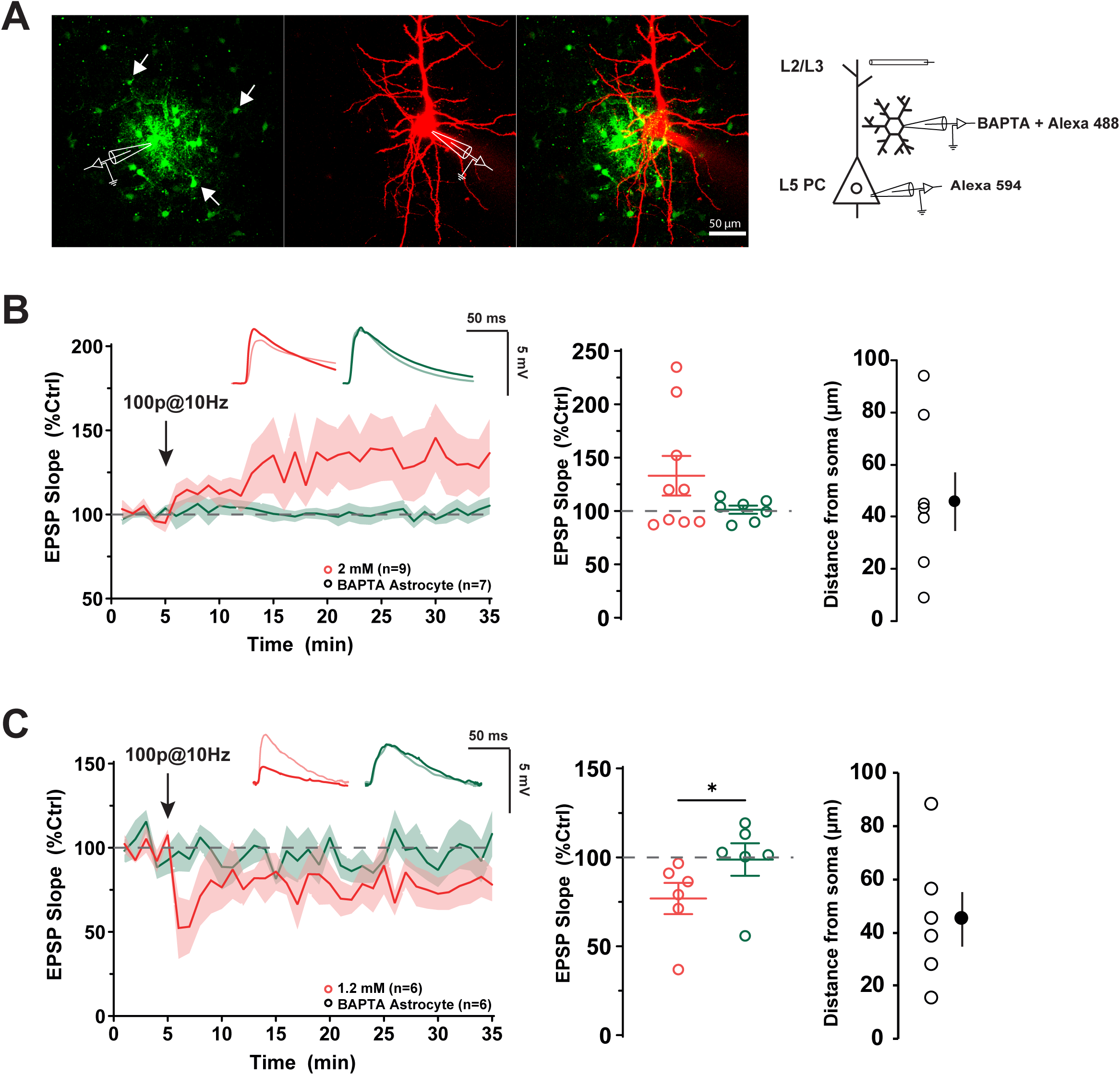
Astrocytes activity modulates t-LTP at L2/L3-L5 synapses in the visual cortex. **A.** Experimental design. Whole-cell recordings were obtained from layer 5 pyramidal neurons, while a stimulating pipette was placed in layer 2/3 to evoke presynaptic stimulation. Simultaneously, a peri-somatic astrocyte was filled with BAPTA (20 mM) for inactivation and Alexa 488 for visualization. **B.** *Left:* Time course of normalized EPSP slope before (light traces) and after (dark traces) pre-post stimulation at 10 Hz in 2 mM condition (red) or during peri-somatic astrocyte network inactivation (green). *Middle:* Quantification of EPSP slope changes in the last 10 minutes of recording, with individual values for each cell represented as red open circles (Control) and green open circles (Astrocytes inactivated). *Right:* Distance of the inactivated astrocytes from the soma of the recorded neuron. **C.** Left: Time course of normalized EPSP slope before (light traces) and after (dark traces) pre-post stimulation at 10 Hz in 1.2 mM condition (red) or during peri-somatic astrocyte network inactivation (green). Middle: Quantification of EPSP slope changes in the last 10 minutes of recording, with individual values for each cell represented as red open circles (Control) and green open circles (Astrocytes inactivated). Right: Distance of the inactivated astrocytes from the soma of the recorded neuron.

### t-LTP at L2/L3-L5 synapses in the visual cortex requires S100β, but not ATP or D-serine

To determine whether S100β contributed to these effects, we repeated the experiment while applying an antibody against S100β (Ab-S100β) during the baseline period of recording and throughout the pre-post protocol at the somatic level of the recorded L5PN. According to our hypothesis, blocking S100β should prevent the decrease in Ca²⁺ and therefore promote t-LTP. However, surprisingly, the application of Ab-S100β shifted t-LTP toward a robust t-LTD (74 % ± 14.2, n = 6, **Fig. 3A**) in 2mM [Ca2+]_e_, and did not impact t-LTD in 1.2 mM [Ca2+]_e_ (63.5 % ± 11.6, n = 5, **Fig. 3B**). As a control, we applied the same antibody after denaturation (20 min at 100°C), which did not affect t-LTP compared to the untreated control condition in 2mM [Ca2+]_e_ (118% ± 10.1, n = 6; **Fig. 3A**). To further ascertain the effects observed with the Ab-S100β, we repeated the experiments in S100βKO mice and found that in both 2mM and 1.2 mM [Ca^2+^]_e_, pre-post protocol at 10 Hz, repeated 100 times, induced t-LTD (64% ± 14.75, n = 6, in 2mM **Fig. 3A**; and 78.1% ± 15.5, n = 5, in 1.2mM **Fig. 3B**). To verify whether these unexpected findings resulted from the release of other gliotransmitters, we assessed the potential involvement of D-serine and ATP in t-LTP by performing recordings in the presence of L701,324, a D-serine/glycine binding site antagonist, and Suramin, a purinergic receptor antagonist (**Fig. 3**). Neither compound prevented the induction of t-LTP (126.3% ± 14.6, n = 5; **Fig. 3C**; 132% ± 22.5, n = 5; **Fig. 3C**), further supporting the critical role of S100β in t-LTP. Together, these results indicate that S100β is essential for normal t-LTP, contradicting our initial hypothesis that its presence would promote t-LTD due to its role as a calcium chelator. The loss of S100β, whether through antibody application or in S100β knockout models, shifts t-LTP toward t-LTD suggesting that its effect on t-LTP may not be entirely mediated by its calcium chelating properties or that they may be location-dependent.

**Figure 3.**
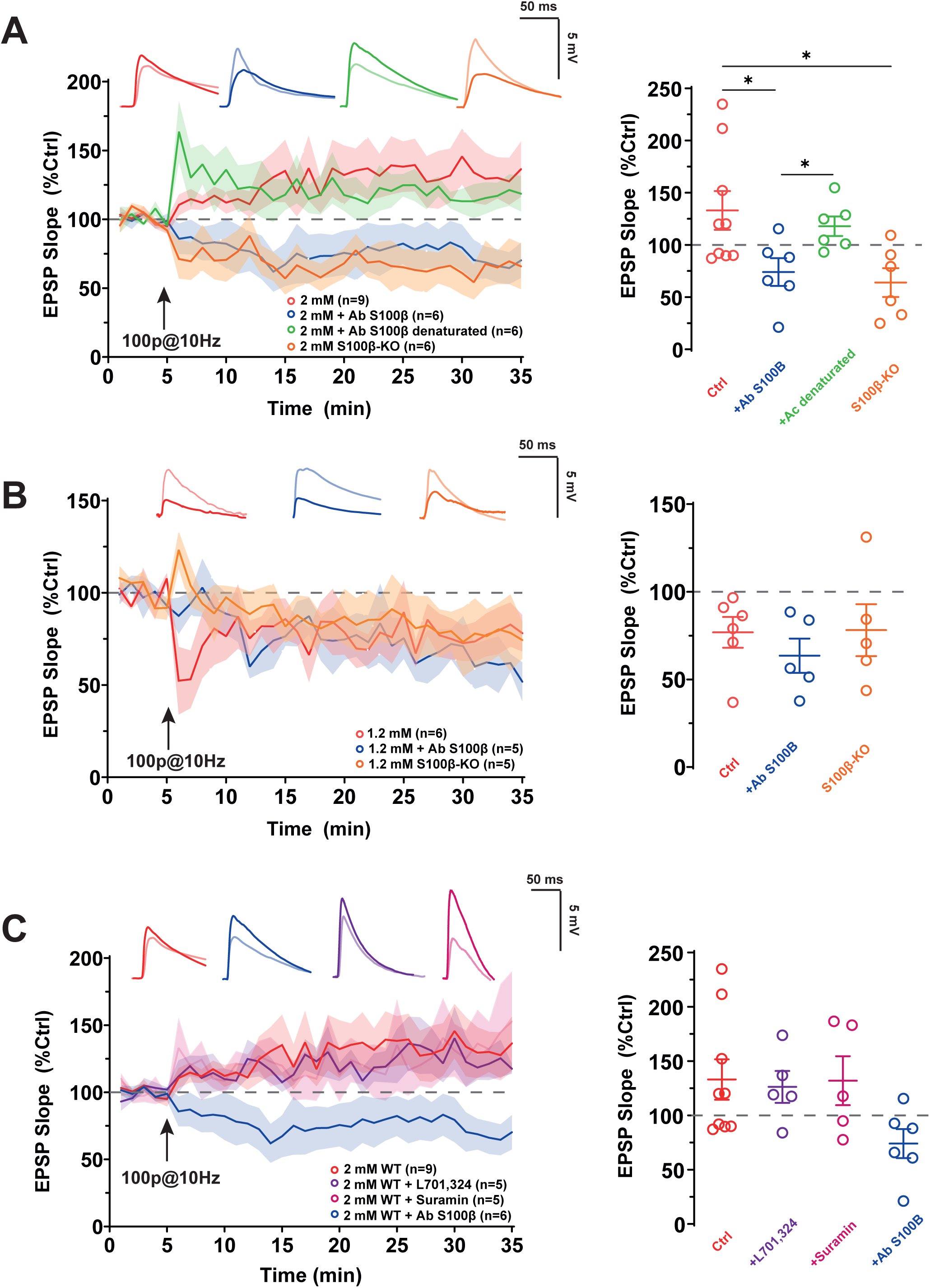
S100β is required for normal t-LTP at L2/L3-L5 synapses in the visual cortex. **A.** *Left:* Time course of normalized EPSP slope before (light traces) and after (dark traces) pre-post stimulation at 10 Hz in the control (red), presence (dark blue) of an antibody against S100β (Ab S100β), the same antibody denatured (green) or in S100βKO mice (orange) in 2 mM extracellular calcium concentration. *Right:* Quantification of EPSP slope changes in the last 10 minutes of recording, with individual values for each cell represented as red open circles (control), dark blue open circles (Ab S100β), green open circles (Ab S100β denaturated) or orange open circles (S100βKO). **B.** *Left:* Time course of normalized EPSP slope before (light traces) and after (dark traces) pre-post stimulation at 10 Hz in the wild-type (red), S100βKO (orange) mice or in presence (dark blue) of an antibody against S100β in S100βKO mice (Ab S100β) in 1.2 mM extracellular calcium concentration. *Right:* Quantification of EPSP slope changes in the last 10 minutes of recording, with individual values for each cell represented as red open circles (WT), orange open circles (S100βKO) and dark blue open circle (S100βKO + Ab S100β). **C.** *Left:* Summary of the time course of normalized EPSP slopes before (light traces) and after (dark traces) 10 Hz pre-post stimulation in WT mice under control conditions (red), or in the presence of L701,324 (purple), Suramin (pink), or anti-S100β antibody (dark blue) in 2mM extracellular calcium concentration. *Right:* Quantification of EPSP slope changes in the last 10 minutes of recording, with individual values for each cell represented as open circles in all the conditions.

### S100β can rescue t-LTP

To further confirm whether S100β is necessary for normal t-LTP at L2/3-L5 synapses in the visual cortex, we repeated the previous experiment in S100βKO mice while locally applying S100β (129 μM) for 5 minutes before the pre-post pairing protocol. This allowed us to fully saturate the local environment without inducing spiking during the protocol itself, which could otherwise alter the results, as STDP is highly sensitive to the number of APs or the resting membrane potential. Confirming the importance of S100β for t-LTP, its application near the somatic compartment in S100βKO mice reversed the observed t-LTD to t-LTP (115 % ± 9, n = 9, **Fig 4A**). Application of S100β in WT mice still induced robust t-LTP but did not further enhance its amplitude compared to WT controls (119% ± 9, n = 6; **Fig. 4B**). Notably, increasing the pairing frequency to 50 Hz, which is known to promote LTP by enhancing postsynaptic calcium influx (Nevian and Sakmann, 2006), was not sufficient to rescue t-LTP in S100βKO mice (73.3% ± 12.7, n = 9; **Fig. 4C**). These results further confirm S100β as a key regulator of t-LTP, though its effect is unlikely mediated by calcium chelation.

**Figure 4.**
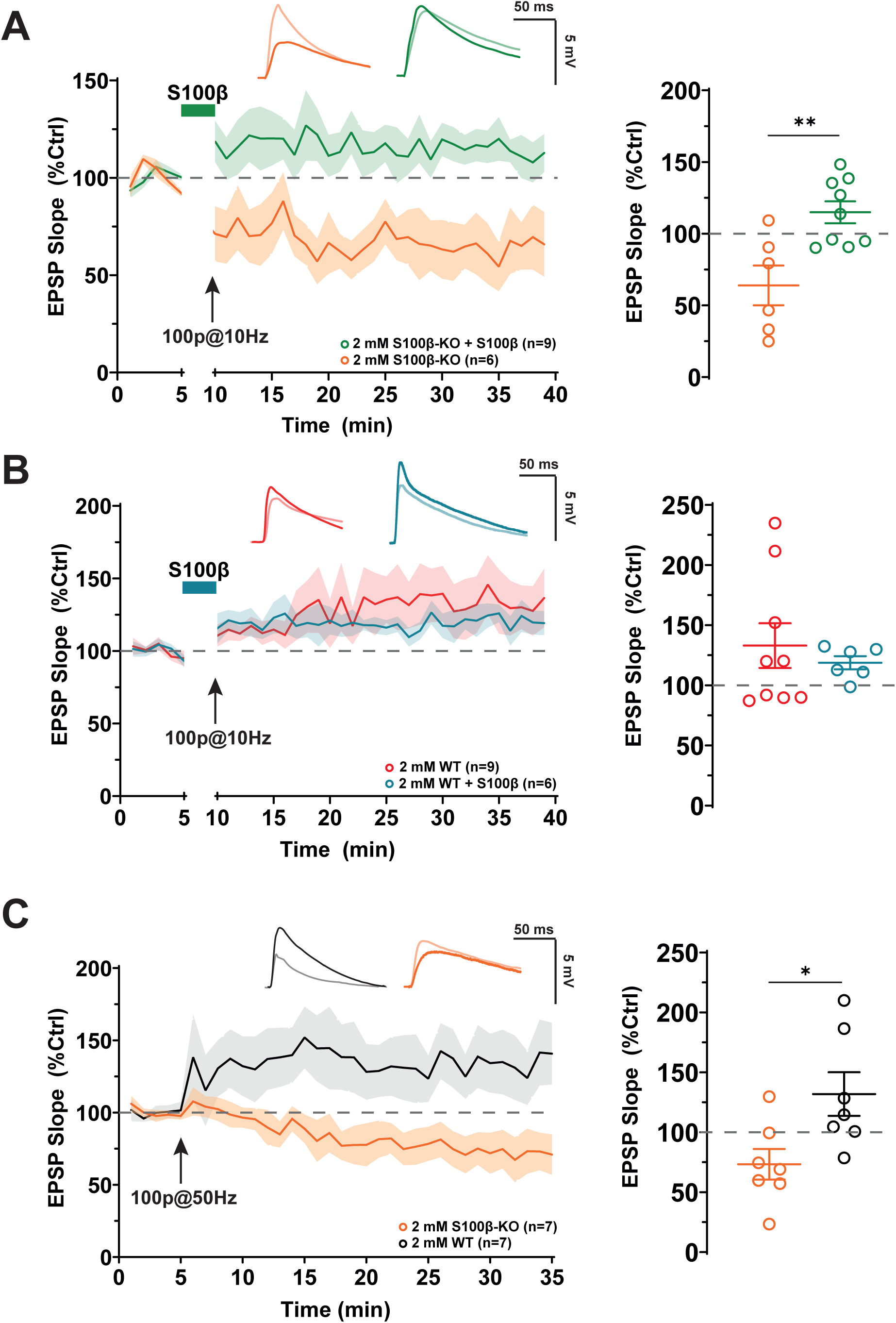
S100β application abolished t-LTD in S100βKO mice. **A.** *Left:* Time course of normalized EPSP slope before (light traces) and after (dark traces) pre-post stimulation at 10 Hz in S100βKO mice in absence (orange) or after a 5min application of the protein S100β (green) before the pairing protocol. *Right:* Quantification of EPSP slope changes in the last 10 minutes of recording, with individual values for each cell represented as orange open circles (S100βKO) and green open circles (S100βKO + S100β). **B**. *Left:* Time course of normalized EPSP slope before (light traces) and after (dark traces) pre-post stimulation at 10 Hz in WT mice in absence (red) or after a 5min application of the protein S100β before the pairing protocol (blue). *Right:* Quantification of EPSP slope changes in the last 10 minutes of recording, with individual values for each cell represented as red open circles (WT) and blue open circles (WT + S100β). **C**. *Left*: Time course of normalized EPSP slope before (light traces) and after (dark traces) pre-post stimulation at 50 Hz in S100βKO mice (orange) or WT (black). *Right*: Quantification of EPSP slope changes in the last 10 minutes of recording, with individual values for each cell represented as orange open circles (S100βKO) and black open circles (WT).

### A-type current inhibition did not restore t-LTP

S100β is a multifaceted protein with numerous known and unknown targets (Donato et al., 2013; Hernández-Ortega et al., 2024). Voltage-gated potassium (K+) channels Kv4.2, which contribute significantly to the somatodendritic A-type current in many neurons have been shown to be inhibited by S100β (Bancroft et al., 2022). These channels attenuate back-propagating action potentials (bAPs) and oppose STDP, which relies on the precise coincidence of EPSPs and bAPs. We therefore hypothesized that the t-LTP-promoting effect of S100β and the t-LTD-promoting effects of its blockade could indirectly result from an effect on these channels. To investigate this possibility, we repeated the experiments in S100βKO mice while blocking Kv4.2 channels using the extracellular application of the Kv4.2 antagonist 4-Aminopyridine (4-AP) or by perfusing an antibody against Kv4.2 (Ab Kv4.2) directly into the recording pipette. However, neither 4-AP (64 % ± 14.7, n = 7, **Fig 5A**) nor intracellular Ab Kv4.2 (66.6 % ± 8, n = 5, **Fig 5B**) restored t-LTP in S100βKO mice, suggesting that the effect of S100β on t-LTP is not mediated by its inhibition of A-type current.

**Figure 5.**
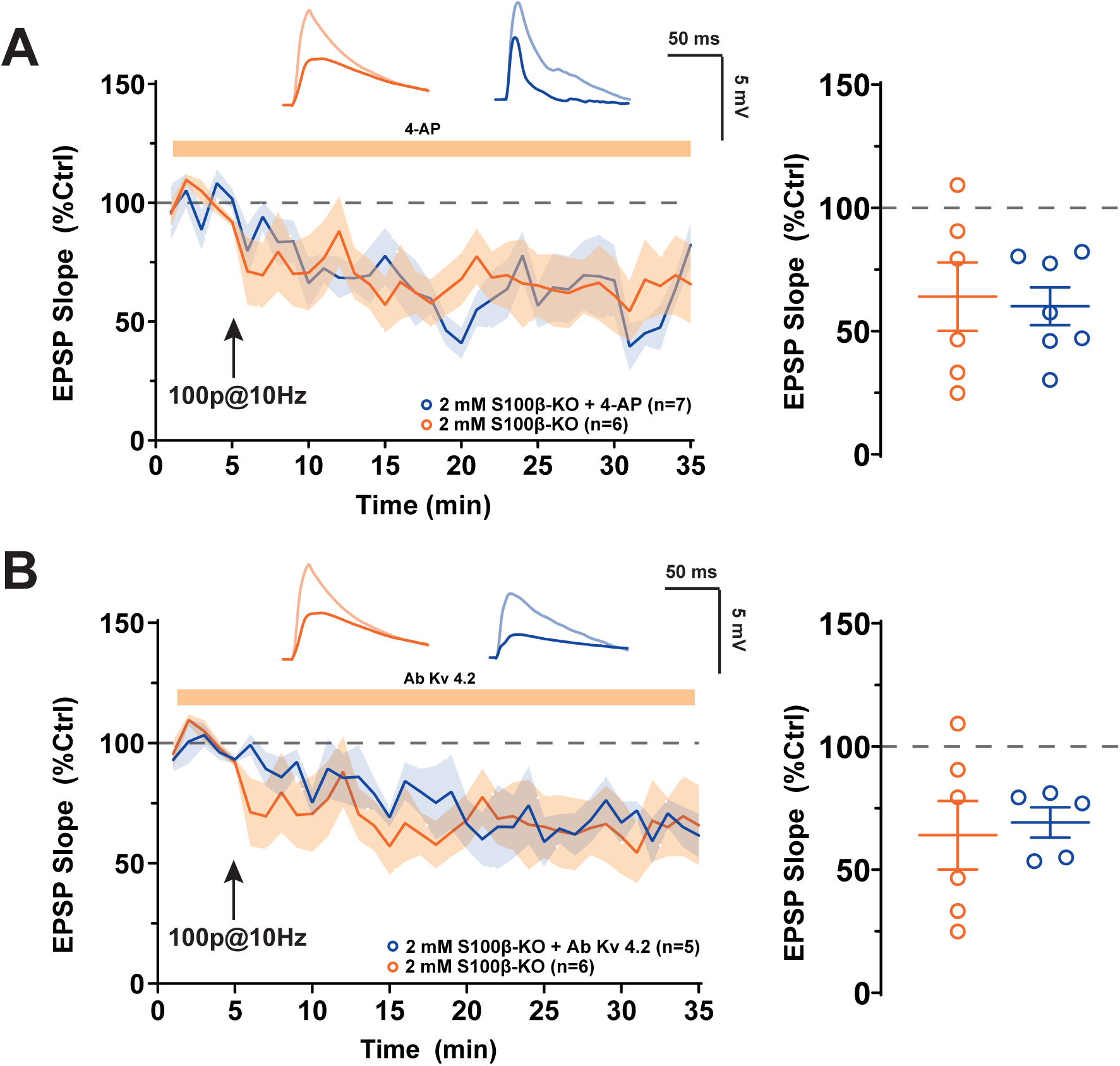
S100β does not exert its effect through Kv 4.2 channel inactivation. **A.** *Left:* Time course of normalized EPSP slope before (light traces) and after (dark traces) pre-post stimulation at 10 Hz in S100βKO mice in absence (orange) or in presence of the Kv 4.2 inhibitor 4-Aminopyridine (blue). *Right:* Quantification of EPSP slope changes in the last 10 minutes of recording, with individual values for each cell represented as orange open circles (S100βKO) and blue open circles (S100βKO + 4-AP). **B.** *Left:* Time course of normalized EPSP slope before (light traces) and after (dark traces) pre-post stimulation at 10 Hz in S100βKO mice in absence (orange) or in presence of an antibody against Kv 4.2 (Ab Kv 4.2) channels in the recording pipette (blue). *Right:* Quantification of EPSP slope changes in the last 10 minutes of recording, with individual values for each cell represented as orange open circles (S100βKO) and blue open circles (S100βKO + Ab Kv 4.2).

## DISCUSSION

### Astrocytes are required for t-LTP at L2/3-L5 synapses of the visual cortex

One of the key findings of our studies is that astrocytic activity is essential for the induction of t-LTP at L2/3-L5 synapses in the visual cortex, as its inactivation prevents this form of plasticity. Our results aligned with previous studies highlighting the critical role of astrocytes in the normal expression of STDP (Sanz-Gálvez et al., 2024). In the barrel cortex, t-LTD has been shown to depend on astrocyte-mediated glutamate release, while in the somatosensory cortex, the release of glutamate and adenosine enables a developmental shift from t-LTD to t-LTP (Min and Nevian, 2012; Martínez-Gallego et al., 2022). Similar mechanisms have been observed in the hippocampus, where astrocytes contribute to t-LTD at CA3-CA1 synapses and in the dentate gyrus, emphasizing their role in synaptic plasticity during development (Falcón-Moya et al., 2020; Cuaya et al., 2024). Furthermore, spontaneous glutamate release from astrocytes has been shown to regulate the t-LTP threshold in the hippocampus (Bonansco et al., 2011). These findings underscore the necessity of incorporating astrocytes into our framework for understanding the cellular mechanisms governing synaptic plasticity and reconsidering existing theories and paradigms. For too long, STDP has been viewed as a relatively simple rule, depending solely on the timing of pre-and postsynaptic activity to modulate synaptic weight, an appealingly elegant model, yet one that has faced criticism regarding its relevance *in vivo*. In addition to other factors known to influence STDP, such as neuromodulation, dendritic spikes, and activity patterns, astrocytes introduce an additional layer of complexity. Among the many factors that regulate STDP astrocytes emerge as powerful modulators through their release of diverse gliotransmitters and their role in ion homeostasis (Araque et al., 2014). However, despite their profound influence, they remain one of the least studied components of synaptic plasticity. One limitation of our study, as well as others, is that it involves the inactivation of a large portion of the astrocytic network, in our case, primarily the peri-somatic astrocytes. However, astrocytes exhibit significant diversity and likely regulate synaptic plasticity in distinct ways depending on their location, whether near the soma, dendrites, or axon. For instance, in the visual cortex, only a subset of astrocytes express S100β (Zhang et al., 2019), suggesting that their role in synaptic plasticity may vary based on their molecular profile, which itself is highly heterogeneous across brain regions (Khakh and Sofroniew, 2015; Endo et al., 2022). Future research on synaptic plasticity should focus on identifying the specific subpopulations of astrocytes involved in its regulation, leveraging recent tools like Astrolight, which enable tagging and targeting of astrocytes engaged in specific activities (Serra et al., 2025).

### Developmental STDP in the visual cortex

Most studies investigating the relationship between astrocytes and STDP have emphasized a key finding: astrocytes play a crucial role in regulating the normal development of STDP across various brain regions (Pérez-Rodríguez et al., 2019; Martínez-Gallego et al., 2022). In the hippocampus and somatosensory cortex, the release of glutamate and adenosine specifically gate a developmental switch from t-LTD to t-LTP. Our study, focusing on the visual cortex of juvenile and young adult animals (P14–P21), deliberately excluded developmental regulation by astrocytes to avoid adding further complexity. However, the visual cortex undergoes significant plastic changes during development, marked by two distinct critical periods: eye opening at P14 and the establishment of binocularity and binocular matching between P21 and P35 (Levelt and Hübener, 2012). In L5 of the visual cortex, astrocytes appear to reach full maturity by P15 but undergo significant changes before this stage (Watanabe et al., 2023). Specifically, gap-junction coupling increases with age, and their arborization becomes denser. This suggests that astrocyte-mediated regulation of STDP in the visual cortex may differ fundamentally before eye opening, especially given that astrocytes have been shown to play a key role in closing the critical period (Ribot et al., 2021). Future research should explore this developmental relationship to determine whether the mechanisms observed in other brain regions also apply to the visual cortex or explore if astrocytes activity is also required during critical periods.

### S100β: a multifaceted protein

This study was originally motivated by two key observations: (1) the dependence of t-LTP on extracellular Ca2+ (Inglebert and Debanne, 2021) and (2) the Ca^2+^-binding properties of S100β, which, when secreted, can significantly alter the local Ca^2+^ environment (Morquette et al., 2015; Ryczko et al., 2021; Gaudel et al., 2025). Surprisingly, however, S100β appears to be more of a prerequisite for normal t-LTP in the visual cortex rather than an inhibitor. Its application promotes t-LTP, while its inhibition favors t-LTD. This highlights the complexity of S100β, which functions beyond regulation of extracellular Ca^2+^ levels. Indeed, S100β has been shown to modulate various ion channels, including Na^+^, K^+^, and Ca^2+^ channels (Hermann et al., 2012). Given the dependence of t-LTP on backpropagating action potentials (bAPs), we initially hypothesized that S100β might promote t-LTP by inhibiting Kv4.2 channels, which act as a brake on bAP propagation (Migliore et al., 1999; Chen et al., 2006; Kim et al., 2007). However, inhibiting Kv4.2 failed to rescue the phenotype observed in S100β knockout (S100βKO) mice, ruling out its involvement. Similarly, NaV 1.6 channels have been shown to facilitate the generation of dendritic spikes, which enhance Ca²⁺ entry into spines and are essential for t-LTP at various synapses (Stuart and Sakmann, 1995; Golding and Spruston, 1998; Kim et al., 2015). Notably, our previous studies have demonstrated that S100β enhances NaV 1.6 channel activity (Morquette et al., 2015; Ryczko et al., 2021), suggesting that this may be one of the mechanisms through which S100β promotes t-LTP. Compared to other members of the S100 protein family, our understanding of S100β targets remains limited. For example, S100A1 has been shown to enhance L-type Ca2+ channel activity (Reppel et al., 2005), facilitate Ca^2+^-induced Ca^2+^ release through ryanodine receptor activation (Wright et al., 2008), and is hypothesized to influence Ca^2+^-activated K^+^ channels, such as SK and BK channels (Kubista et al., 1999), which are involved in regulating EPSP amplitude and STDP induction (Jones et al., 2017; Tazerart et al., 2022). Additionally, S100 proteins directly interact with receptors known to modulate STDP, including dopamine D2 metabotropic receptors, which are implicated in promoting t-LTD (Liu et al., 2008).

### S100β in other forms of plasticity

Synaptic plasticity is not the only mechanism underlying memory formation; other processes, such as the modification of intrinsic plasticity, also play a role (Debanne et al., 2019). This latter process often involves the regulation of ion channels, which can be influenced indirectly or directly by the effects of S100 family proteins like S100β. These regulatory changes can impact various aspects of neuronal excitability, including resting membrane potential, EPSP integration, and action potential (AP) threshold (Debanne and Russier, 2019). Reduction in extracellular Ca^2+^ concentration is directly linked to an increase in excitability (Forsberg et al., 2019, 2025) through an effect on NaV channels. The same effect is observed during S100β application, which reduce [Ca^2+^]_e_, and potentiate the persistent sodium current (I_NaP_) carried by NaV 1.6, lowering AP threshold (Morquette et al., 2015; Ryczko et al., 2021; Gaudel et al., 2025). All the ion channels mentioned earlier play a role in regulating intrinsic excitability, positioning S100β as a strong candidate for modulating this process. For example, the long-term potentiation of intrinsic excitability (LTP-IE) observed in the visual cortex required Protein Kinase A activity which is a direct target of S100 family protein (Cudmore and Turrigiano, 2004; Reppel et al., 2005). While most studies have focused on the role of astrocytes in synaptic plasticity, including STDP and other forms, their involvement in intrinsic plasticity remains an area worth exploring.

## CONCLUSION

Our findings at L2/3-L5 synapses in the visual cortex build upon previous studies in the hippocampus and somatosensory cortex, which have identified astrocyte activity as a prerequisite for normal t-LTP, t-LTD, or both. Our study specifically highlights the role of S100β in this regulatory process, an aspect that has received considerably less attention compared to other gliotransmitters such as adenosine or glutamate.

Moreover, our results underscore how little is known about S100β’s molecular targets and physiological functions beyond its calcium-binding role. Given its potential to modulate various forms of synaptic plasticity through its effects on ion channels and Ca²⁺-dependent processes, S100β may represent a crucial regulator of neuronal plasticity. Future studies will undoubtedly shed light on the many unanswered questions raised by this work.

## METHODS

### Animals

All experiments were conducted in accordance with the rules of the Canadian Institutes of Health Research and were approved by the Animal Care and Use Committee of the University of Montreal (Protocol #23-207). C57BL/6J (WT) and S100β-Knockout (S100β-KO) transgenic mice (B6Brd; B6N-Tyrc-Brd S100btm1a(EUCOMM)Wtsi/WtsiCnbc, S100b (MCFR; EPD0157_6_C08) Wellcome Trust Sanger Institute) were used in this study. WT or S100βKO were fed ad libitum and housed with a 12 h light/dark cycle. WT and S100βKO male and female mice were used for experiments.

### Electrophysiology

Visual Cortex *ex vivo* slices were obtained from P14-P21 old WT or S100βKO mice. Mice were deeply anesthetized with isoflurane and killed by decapitation. Coronal Slices (350 µm) were cut on a vibratome (Leica Microsystems, VT1000S) in an ice-cold (0-4°C) sucrose-based solution containing the following (in mM): 219 sucrose, 26 NaHCO_3_, 10 dextrose, 3 KCl, 0.2 CaCl_2_, 1.25 KH_2_PO_4_ and 4 MgSO_4_ (pH 7.3-7.4, 300-320 mOsmol/kg) and were transferred at 32°C in regular ACSF containing the following (in mM): 124 NaCl, 3 KCl, 1.3 MgSO_4_, 26 NaHCO_3_, 2 CaCl_2_ (or 1.2), 1.25 KH_2_PO_4_ and 10 dextrose saturated with 95% O2/5% CO2 (pH 7.3-7.4, 300-320 mOsmol/kg) for 15 min before resting at room temperature (RT) for 1 h in oxygenated (95% O2/5% CO2) ACSF. Recordings were obtained using an Axopatch 200B or Multiclamp 700B (Molecular Devices) amplifier and pClamp10.4 software. Data were sampled at 33 kHz, filtered at 3 kHz, and digitized by a Digidata 1440A (Molecular Devices. Excitatory postsynaptic potentials (EPSPS) were elicited in the layer 2/3 of the visual cortex by using glass microelectrodes filled with ACSF. EPSPs were elicited at 0.1 Hz by a digital stimulator that was fed by a stimulation isolator unit (A320, World Precision Instruments). All data analyses were performed with custom-written software in Igor Pro 8 (Wavemetrics) or with Clampfit (Molecular Devices). The EPSP slope was measured as an index of synaptic strength. Access resistance was monitored throughout the recording and only experiments with stable resistance were kept (changes <20%). Recordings were made from layer 5 pyramidal neurons, electrodes were filled with a solution containing the following (in mM): 140 K-gluconate, 5 NaCl, 20 KCl, 10 HEPES, 2 MgCl_2_, 0.5 EGTA, 0.4 Tris-GTP and 2 Tris-ATP. In experiments where astrocytes were inactivated, BAPTA (20 mM) was added to the internal solution of the electrode used to patch the astrocyte, and recordings were initiated only after a 10-minute period to ensure adequate diffusion of BAPTA throughout the astrocytic network

### Drugs

The following drugs were locally applied near the recorded cells using positive pressure pulses (Picospritzer III): Monoclonal anti-S100β antibodies (mouse anti-S100β, Sigma Aldrich #S2532) and S100β (Produced in the Department of Biochemistry of the Universite de Montreal as described in Morquette et al., 2015). 4-Aminopyridine (100 μM, Tocris #0940), Suramin (100 μM, Tocris #1472), and L701,324 (100 μM, Tocris #0907), were bath applied, and antibody against Kv 4.2 (1 μg/mL, Alomone labs #APC-023) were used in the recoding pipette as described in Lee et al., 2023.

### Data and analysis

Data are presented as mean ± standard error to the mean (SEM). Statistical differences between experimental conditions were calculated, not assuming Gaussian distributions or equal variances, using the Mann–Whitney U test or Wilcoxon rank-signed test. Statistically significant differences were established at *P < 0.05, **P < 0.01 and ***P < 0.001; NS indicates not significant. Data analysis was performed using SPSS (IBM SPSS Statistics).

## DECLARATIONS

### Author contributions

R.S.G., Y.I and A.K wrote the paper. R.S.G and Y.I collected and analyzed the data and built the figures. All authors contributed to the article and approved the submitted version.

## Acknowledgements

We thank Dr. Dorly Verdier and Dr. Fanny Gaudel for critically reading a preliminary version of the manuscript and members of A.K laboratories for comments on the manuscript. We are especially grateful to our esteemed colleagues, Dr. James G. Omichinski and Dr. Haytham Wahba, from the Department of Biochemistry at the Université de Montréal, for their invaluable contribution to the production of the S100β protein. Figure 1 was partially created using an image from NIAID Visual & Medical Arts (https://bioart.niaid.nih.gov/bioart/370), a free illustration collection from the NIH.

## Availability of data and materials

All data generated or analysed during this study are included in this published article.

## Preprint

This manuscript was posted on a preprint.

## REFERENCES

Araque, A., Carmignoto, G., Haydon, P. G., Oliet, S. H. R., Robitaille, R., and Volterra, A. (2014). Gliotransmitters Travel in Time and Space. Neuron 81, 728–739. doi: 10.1016/j.neuron.2014.02.007

Araque, A., Parpura, V., Sanzgiri, R. P., and Haydon, P. G. (1998). Glutamate-dependent astrocyte modulation of synaptic transmission between cultured hippocampal neurons. European Journal of Neuroscience 10, 2129–2142. doi: 10.1046/j.1460-9568.1998.00221.x

Bancroft, E. A., De La Mora, M., Pandey, G., Zarate, S. M., and Srinivasan, R. (2022). Extracellular S100B inhibits A-type voltage-gated potassium currents and increases L-type voltage-gated calcium channel activity in dopaminergic neurons. Glia 70, 2330–2347. doi: 10.1002/glia.24254

Bi, G., and Poo, M. (1998). Synaptic Modifications in Cultured Hippocampal Neurons: Dependence on Spike Timing, Synaptic Strength, and Postsynaptic Cell Type. J Neurosci 18, 10464–10472. doi: 10.1523/JNEUROSCI.18-24-10464.1998

Bonansco, C., Couve, A., Perea, G., Ferradas, C. Á., Roncagliolo, M., and Fuenzalida, M. (2011). Glutamate released spontaneously from astrocytes sets the threshold for synaptic plasticity. European Journal of Neuroscience 33, 1483–1492. doi: 10.1111/j.1460-9568.2011.07631.x

Brzosko, Z., Mierau, S. B., and Paulsen, O. (2019). Neuromodulation of Spike-Timing-Dependent Plasticity: Past, Present, and Future. Neuron 103, 563–581. doi: 10.1016/j.neuron.2019.05.041

Chen, X., Yuan, L.-L., Zhao, C., Birnbaum, S. G., Frick, A., Jung, W. E., et al. (2006). Deletion of Kv4.2 gene eliminates dendritic A-type K+ current and enhances induction of long-term potentiation in hippocampal CA1 pyramidal neurons. J Neurosci 26, 12143–12151. doi: 10.1523/JNEUROSCI.2667-06.2006

Chindemi, G., Abdellah, M., Amsalem, O., Benavides-Piccione, R., Delattre, V., Doron, M., et al. (2022). A calcium-based plasticity model for predicting long-term potentiation and depression in the neocortex. Nat Commun 13, 3038. doi: 10.1038/s41467-022-30214-w

Cuaya, H. C., Martínez-Gallego, I., and Rodríguez-Moreno, A. (2024). Astrocytes mediate two forms of spike timing-dependent depression at entorhinal cortex-hippocampal synapses. eLife 13. doi: 10.7554/eLife.98031.1

Cudmore, R. H., and Turrigiano, G. G. (2004). Long-Term Potentiation of Intrinsic Excitability in LV Visual Cortical Neurons. Journal of Neurophysiology 92, 341–348. doi: 10.1152/jn.01059.2003

Debanne, D., Gähwiler, B. H., and Thompson, S. M. (1996). Cooperative interactions in the induction of long-term potentiation and depression of synaptic excitation between hippocampal CA3-CA1 cell pairs in vitro. Proc. Natl. Acad. Sci. U.S.A. 93, 11225–11230. doi: 10.1073/pnas.93.20.11225

Debanne, D., and Inglebert, Y. (2023). Spike timing-dependent plasticity and memory. Current Opinion in Neurobiology 80, 102707. doi: 10.1016/j.conb.2023.102707

Debanne, D., Inglebert, Y., and Russier, M. (2019). Plasticity of intrinsic neuronal excitability. Current Opinion in Neurobiology 54, 73–82. doi: 10.1016/j.conb.2018.09.001

Debanne, D., and Russier, M. (2019). The contribution of ion channels in input-output plasticity. Neurobiology of Learning and Memory 166, 107095. doi: 10.1016/j.nlm.2019.107095

Donato, R., Cannon, B. R., Sorci, G., Riuzzi, F., Hsu, K., Weber, D. J., et al. (2013). Functions of S100 Proteins. Curr Mol Med 13, 24–57.

Endo, F., Kasai, A., Soto, J. S., Yu, X., Qu, Z., Hashimoto, H., et al. (2022). Molecular basis of astrocyte diversity and morphology across the CNS in health and disease. Science 378, eadc9020. doi: 10.1126/science.adc9020

Falcón-Moya, R., Pérez-Rodríguez, M., Prius-Mengual, J., Andrade-Talavera, Y., Arroyo-García, L. E., Pérez-Artés, R., et al. (2020). Astrocyte-mediated switch in spike timing-dependent plasticity during hippocampal development. Nat Commun 11, 4388. doi: 10.1038/s41467-020-18024-4

Feldman, D. E. (2012). The Spike-Timing Dependence of Plasticity. Neuron 75, 556–571. doi: 10.1016/j.neuron.2012.08.001

Forsberg, M., Seth, H., Björefeldt, A., Lyckenvik, T., Andersson, M., Wasling, P., et al. (2019). Ionized calcium in human cerebrospinal fluid and its influence on intrinsic and synaptic excitability of hippocampal pyramidal neurons in the rat. Journal of Neurochemistry 149, 452–470. doi: 10.1111/jnc.14693

Forsberg, M., Zhou, D., Jalali, S., Faravelli, G., Seth, H., Björefeldt, A., et al. (2025). Evaluation of mechanisms involved in regulation of intrinsic excitability by extracellular calcium in CA1 pyramidal neurons of rat. Journal of Neurochemistry 169, e16209. doi: 10.1111/jnc.16209

Gaudel, F., Giraud, J., Morquette, P., Couillard-Larocque, M., Verdier, D., and Kolta, A. (2025). Astrocyte-induced firing in primary afferent axons. iScience 0. doi: 10.1016/j.isci.2025.112006

Golding, N. L., and Spruston, N. (1998). Dendritic sodium spikes are variable triggers of axonal action potentials in hippocampal CA1 pyramidal neurons. Neuron 21, 1189–1200. doi: 10.1016/s0896-6273(00)80635-2

Henneberger, C., Papouin, T., Oliet, S. H. R., and Rusakov, D. A. (2010). Long-term potentiation depends on release of d-serine from astrocytes. Nature 463, 232–236. doi: 10.1038/nature08673

Hermann, A., Donato, R., Weiger, T. M., and Chazin, W. J. (2012). S100 Calcium Binding Proteins and Ion Channels. Front Pharmacol 3, 67. doi: 10.3389/fphar.2012.00067

Hernández-Ortega, K., Canul-Euan, A. A., Solis-Paredes, J. M., Borboa-Olivares, H., Reyes-Muñoz, E., Estrada-Gutierrez, G., et al. (2024). S100B actions on glial and neuronal cells in the developing brain: an overview. Front. Neurosci. 18. doi: 10.3389/fnins.2024.1425525

Huang, S., Huganir, R. L., and Kirkwood, A. (2013). Adrenergic Gating of Hebbian Spike-Timing-Dependent Plasticity in Cortical Interneurons. Journal of Neuroscience 33, 13171–13178. doi: 10.1523/JNEUROSCI.5741-12.2013

Inglebert, Y., Aljadeff, J., Brunel, N., and Debanne, D. (2020). Synaptic plasticity rules with physiological calcium levels. Proc. Natl. Acad. Sci. U.S.A. 117, 33639–33648. doi: 10.1073/pnas.2013663117

Inglebert, Y., and Debanne, D. (2021). Calcium and Spike Timing-Dependent Plasticity. Front. Cell. Neurosci. 15, 727336. doi: 10.3389/fncel.2021.727336

Jones, S. L., To, M.-S., and Stuart, G. J. (2017). Dendritic small conductance calcium-activated potassium channels activated by action potentials suppress EPSPs and gate spike-timing dependent synaptic plasticity. eLife 6, e30333. doi: 10.7554/eLife.30333

Khakh, B. S., and Sofroniew, M. V. (2015). Diversity of astrocyte functions and phenotypes in neural circuits. Nat Neurosci 18, 942–952. doi: 10.1038/nn.4043

Kim, J., Jung, S.-C., Clemens, A. M., Petralia, R. S., and Hoffman, D. A. (2007). Regulation of dendritic excitability by activity-dependent trafficking of the A-type K+ channel subunit Kv4.2 in hippocampal neurons. Neuron 54, 933–947. doi: 10.1016/j.neuron.2007.05.026

Kim, Y., Hsu, C.-L., Cembrowski, M. S., Mensh, B. D., and Spruston, N. (2015). Dendritic sodium spikes are required for long-term potentiation at distal synapses on hippocampal pyramidal neurons. Elife 4, e06414. doi: 10.7554/eLife.06414

Kubista, H., Donato, R., and Hermann, A. (1999). S100 calcium binding protein affects neuronal electrical discharge activity by modulation of potassium currents. Neuroscience 90, 493–508. doi: 10.1016/s0306-4522(98)00422-9

Lee, S. Y., Kwon, M., Ho, W. K., and Lee, S.-H. (2023). Kv4.2 regulates baseline synaptic strength by inhibiting R-type channel-mediated calcium signaling in the hippocampus. 2023.12.05.570317. doi: 10.1101/2023.12.05.570317

Levelt, C. N., and Hübener, M. (2012). Critical-Period Plasticity in the Visual Cortex. Annual Review of Neuroscience 35, 309–330. doi: 10.1146/annurev-neuro-061010-113813

Liu, Y., Buck, D. C., and Neve, K. A. (2008). Novel interaction of the dopamine D2 receptor and the Ca2+ binding protein S100B: role in D2 receptor function. Mol Pharmacol 74, 371–378. doi: 10.1124/mol.108.044925

Martínez-Gallego, I., Pérez-Rodríguez, M., Coatl-Cuaya, H., Flores, G., and Rodríguez-Moreno, A. (2022). Adenosine and Astrocytes Determine the Developmental Dynamics of Spike Timing-Dependent Plasticity in the Somatosensory Cortex. J. Neurosci. 42, 6038–6052. doi: 10.1523/JNEUROSCI.0115-22.2022

McFarlan, A. R., Chou, C. Y. C., Watanabe, A., Cherepacha, N., Haddad, M., Owens, H., et al. (2023). The plasticitome of cortical interneurons. Nat Rev Neurosci 24, 80–97. doi: 10.1038/s41583-022-00663-9

Migliore, M., Hoffman, D. A., Magee, J. C., and Johnston, D. (1999). Role of an A-Type K+ Conductance in the Back-Propagation of Action Potentials in the Dendrites of Hippocampal Pyramidal Neurons. J Comput Neurosci 7, 5–15. doi: 10.1023/A:1008906225285

Min, R., and Nevian, T. (2012). Astrocyte signaling controls spike timing–dependent depression at neocortical synapses. Nat Neurosci 15, 746–753. doi: 10.1038/nn.3075

Morquette, P., Verdier, D., Kadala, A., Féthière, J., Philippe, A. G., Robitaille, R., et al. (2015). An astrocyte-dependent mechanism for neuronal rhythmogenesis. Nat Neurosci 18, 844–854. doi: 10.1038/nn.4013

Nevian, T., and Sakmann, B. (2006). Spine Ca2+ signaling in spike-timing-dependent plasticity. J Neurosci 26, 11001–11013. doi: 10.1523/JNEUROSCI.1749-06.2006

Pérez-Rodríguez, M., Arroyo-García, L. E., Prius-Mengual, J., Andrade-Talavera, Y., Armengol, J. A., Pérez-Villegas, E. M., et al. (2019). Adenosine Receptor-Mediated Developmental Loss of Spike Timing-Dependent Depression in the Hippocampus. Cereb Cortex 29, 3266–3281. doi: 10.1093/cercor/bhy194

Pike, F. G., Meredith, R. M., Olding, A. W. A., and Paulsen, O. (1999). Postsynaptic bursting is essential for ‘Hebbian’ induction of associative long-term potentiation at excitatory synapses in rat hippocampus. The Journal of Physiology 518, 571–576. doi: 10.1111/j.1469-7793.1999.0571p.x

Reppel, M., Sasse, P., Piekorz, R., Tang, M., Roell, W., Duan, Y., et al. (2005). S100A1 enhances the L-type Ca2+ current in embryonic mouse and neonatal rat ventricular cardiomyocytes. J Biol Chem 280, 36019–36028. doi: 10.1074/jbc.M504750200

Ribot, J., Breton, R., Calvo, C.-F., Moulard, J., Ezan, P., Zapata, J., et al. (2021). Astrocytes close the mouse critical period for visual plasticity. Science 373, 77–81. doi: 10.1126/science.abf5273

Ryczko, D., Hanini-Daoud, M., Condamine, S., Bréant, B. J. B., Fougère, M., Araya, R., et al. (2021). S100β-mediated astroglial control of firing and input processing in layer 5 pyramidal neurons of the mouse visual cortex. The Journal of Physiology 599, 677–707. doi: 10.1113/JP280501

Sanz-Gálvez, R., Falardeau, D., Kolta, A., and Inglebert, Y. (2024). The role of astrocytes from synaptic to non-synaptic plasticity. Front. Cell. Neurosci. 18. doi: 10.3389/fncel.2024.1477985

Serra, I., Martín-Monteagudo, C., Sánchez Romero, J., Quintanilla, J. P., Ganchala, D., Arevalo, M.-A., et al. (2025). Astrocyte ensembles manipulated with AstroLight tune cue-motivated behavior. Nat Neurosci, 1–11. doi: 10.1038/s41593-025-01870-0

Stuart, G., and Sakmann, B. (1995). Amplification of EPSPs by axosomatic sodium channels in neocortical pyramidal neurons. Neuron 15, 1065–1076. doi: 10.1016/0896-6273(95)90095-0

Tan, Z., Liu, Y., Xi, W., Lou, H., Zhu, L., Guo, Z., et al. (2017). Glia-derived ATP inversely regulates excitability of pyramidal and CCK-positive neurons. Nat Commun 8, 13772. doi: 10.1038/ncomms13772

Tazerart, S., Blanchard, M. G., Miranda-Rottmann, S., Mitchell, D. E., Navea Pina, B., Thomas, C. I., et al. (2022). Selective activation of BK channels in small-headed dendritic spines suppresses excitatory postsynaptic potentials. The Journal of Physiology 600, 2165–2187. doi: 10.1113/JP282303

Watanabe, A., Guo, C., and Sjöström, P. J. (2023). The developmental profile of visual cortex astrocytes. iScience 26, 106828. doi: 10.1016/j.isci.2023.106828

Wright, N. T., Prosser, B. L., Varney, K. M., Zimmer, D. B., Schneider, M. F., and Weber, D. J. (2008). S100A1 and calmodulin compete for the same binding site on ryanodine receptor. J Biol Chem 283, 26676– 26683. doi: 10.1074/jbc.M804432200

Zhang, J.-C., Lau, P.-M., and Bi, G.-Q. (2009). Gain in sensitivity and loss in temporal contrast of STDP by dopaminergic modulation at hippocampal synapses. Proc. Natl. Acad. Sci. U.S.A. 106, 13028– 13033. doi: 10.1073/pnas.0900546106

Zhang, Z., Ma, Z., Zou, W., Guo, H., Liu, M., Ma, Y., et al. (2019). The Appropriate Marker for Astrocytes: Comparing the Distribution and Expression of Three Astrocytic Markers in Different Mouse Cerebral Regions. BioMed Research International 2019, 9605265. doi: 10.1155/2019/9605265

